# Precuneus stimulation alters the neural dynamics of autobiographical memory retrieval

**DOI:** 10.1101/592436

**Authors:** Melissa Hebscher, Christine Ibrahim, Asaf Gilboa

## Abstract

Autobiographical memory (AM) unfolds over time, but little is known about the dynamics of its retrieval. Space-based models of memory implicate the hippocampus, retrosplenial cortex, and precuneus in early memory computations. Here we used inhibitory continuous theta burst stimulation (cTBS) to determine the causal role of the precuneus in the temporal dynamics of AM retrieval. Compared to vertex, precuneus stimulation altered evoked neural activity during early memory construction, as early as 400 ms after cue presentation, as well as during later memory elaboration. We further identified a parietal late positive component during memory elaboration, the amplitude of which was associated with spatial perspective during recollection. Precuneus stimulation disrupted this association, suggesting that this region plays a crucial role in the neural representation of spatial perspective during AM. These findings help clarify the neural correlates of early memory retrieval and demonstrate a causal role for the precuneus in the temporal dynamics of AM retrieval.

## 1. Introduction

Autobiographical memories (AMs) unfold over time, allowing individuals to re-experience events from their past in detail, often embedded in a spatial context. While a rich body of literature has examined the neural correlates of AM, few studies to date have studied the temporal dynamics of its retrieval. Predictions about the temporal dynamics of AM may be gleaned from space-based theories and models of memory. One such model argues that transformations between spatial perspectives during early memory retrieval are crucial for spatial and episodic memory (Becker & Burgess, 2001; Byrne, Becker, & Burgess, 2007). According to this model, the hippocampus is important for third-person, allocentric space-based representations during early memory retrieval. The retrosplenial cortex translates these into first-person, egocentric representations, which are then used by the precuneus for visual imagery (Bicanski & Burgess, 2018; Byrne et al., 2007). Together these models implicate the hippocampus, retrosplenial cortex, and precuneus in early space-based computations which are thought to be important for memory retrieval. While prior studies have implicated these regions in AM more generally (Svoboda, McKinnon, & Levine, 2006 for review), and in perspective-taking during AM specifically (Freton et al., 2014; Hebscher, Levine, & Gilboa, 2018; Jacques, Szpunar, & Schacter, 2016), little evidence exists on the temporal dynamics of their recruitment.

In contrast to the extensive literature on episodic AM using functional magnetic resonance imaging (fMRI), only a small number of studies have used electrophysiological recordings to study AM retrieval dynamics. Early studies used electroencephalography (EEG) to measure slow cortical potentials during AM retrieval and identified early left frontal negativity and later temporal and occipital negativity (Conway, Pleydell-Pearce, & Whitecross, 2001; Conway, Pleydell-Pearce, Whitecross, & Sharpe, 2003). More recently, Renoult et al (2016) showed that autobiographically significant names (famous names that easily bring to mind personal episodic memories) are associated with increased amplitude of a signature of episodic recollection, the late positive component (LPC), compared to names with low autobiographical significance. The LPC is typically largest over posterior electrodes and is thought to reflect recollection-sensitive activity in parietal and medial temporal lobes (MTL) (Addante, Ranganath, Olichney, & Yonelinas, 2012; Düzel, Vargha-Khadem, Heinze, & Mishkin, 2001; Hoppstädter, Baeuchl, Diener, Flor, & Meyer, 2015; Rugg & Curran, 2007). While the LPC has been studied extensively in lab-based episodic memory tasks and may be sensitive to autobiographical significance (Renoult et al., 2016), it has never been directly studied in the context of AM retrieval.

In the present study we combined inhibitory transcranial magnetic stimulation (TMS) with electrophysiological measures using magnetoencephalography (MEG) to determine the causal role of the precuneus in rapid dynamic AM processes. Previous studies have used TMS to understand the causal role of parietal regions in autobiographical (Bonnici, Cheke, Green, FitzGerald, & Simons, 2018; Hebscher, Meltzer, & Gilboa, 2019; Thakral, Madore, & Schacter, 2017) and episodic memory (Bonnì et al., 2015; Nilakantan et al., 2017; Wang et al., 2014; Wang & Voss, 2015; Yazar et al., 2014), some of which have shown associated and sustained alterations of neural activity (Hebscher et al., 2019; Nilakantan et al., 2017; Wang et al., 2014; Wang & Voss, 2015). We aimed to observe how altering activity in the precuneus affects the temporal dynamics of AM retrieval at the behavioural level by measuring reaction times and early spatial representations, and at the neural level using MEG. Given the proposed early role of the precuneus in the temporal dynamics of AM (Byrne, Becker, & Burgess, 2007), we hypothesized that inhibitory precuneus stimulation would influence retrieval dynamics during both early search (access) and elaboration (re-experiencing) phases of AM retrieval. Specifically, we hypothesized that (i) stimulation would disrupt the dynamics of memory retrieval as reflected by behaviour (reaction times and ratings); (ii) stimulation would disrupt early components of the event related fields (ERFs); and (iii) ERF disruption would be related to behavioural measures of spatial processing. One specific component we were interested in was the LPC, which peaks around 600-800 ms during lab-based episodic memory (Rugg & Curran, 2007). We examined the LPC and its behavioural correlates to determine whether this component reflects recollection during early stages of AM re-experiencing, as it does in lab-based episodic memory tasks.

## 2. Methods

### 2.1. Participants

23 healthy young participants (14 females; mean age = 26.3, range = 19-36) were tested on a within-subjects combined TMS-MEG paradigm. Participants were recruited from the Rotman Research Institute’s healthy volunteer pool. Participants had completed an average of 16.4 (range = 14-21) years of formal education, were all right-handed, native or fluent English speakers, had normal or corrected-to-normal vision, and were free from a history of neurological illness or injury, psychiatric condition, substance abuse, or serious medical conditions. Based on TMS safety guidelines, participants were excluded if they had a history of losing consciousness (fainting), had a prior experience of a seizure, or had a diagnosis or family history of epilepsy. All participants provided informed consent prior to participating in the experiment in accordance with the Rotman Research Institute/Baycrest Hospital ethical guidelines.

### 2.2. Procedure

Participants received cTBS to their left precuneus and to a control region (vertex) on separate days, at least 24 hours apart (mean = 5.4 days, SD = 5.3 days). Immediately following cTBS, participants completed an AM task inside the MEG scanner which was located nearby. All participants completed the MEG scan within an average of 27.4 minutes (SD = 3.8) measured from the end of cTBS. Average time between the end of cTBS and the start of MEG was 6.04 minutes (SD = 1.5). Anatomical MRIs for each participant were collected in a separate session and used for identifying stimulation locations (see below).

#### 2.2.1. Stimuli and task

At least 48 hours prior to the study, participants provided the names of familiar places, objects and people in an online interview. Locations, people and objects were used as cues because they are elements that commonly make up an event (Addis, Pan, Vu, Laiser, & Schacter, 2009; Burgess, Maguire, et al., 2001). Participants were instructed to name the first twenty items that came to mind, to limit items to those encountered within the past year, and to limit personal objects to those that are not tied to a particular location.

Based on the online interview, sixty cue words were created for each participant, 20 per category. These were randomly divided between the two stimulation sessions so that each session included 30 cues, and each session was broken down into 3 runs to be used in the MEG scanner. E-Prime 1.2 software was used to display the items and collect response data. Items were presented in a randomized order. Participants were instructed to use the words as cues in order to recall personal events that are specific in time that had occurred in approximately the last year, not including the past week. Specific events were defined as “past events from a specific time and place for which you were personally involved.” Cue words were displayed for a maximum of 10 s and participants were instructed to retrieve a specific past event related to the cue as quickly as possible. Participants were asked to press a button on the response box corresponding to their right index finger as soon as a memory came to mind. Trials in which no memory was retrieved were discarded. The retrieval phase was terminated when a memory was retrieved, after which participants saw a slide asking “What was the very first thing that came to mind”, and had to choose one of the following four options: person, object, place, other. An elaboration phase followed in which participants were prompted to imagine the event in as much detail as possible for 8 s. Next, participants rated the memory on four scales aimed at measuring different phenomenological characteristics of the memory. They were given a maximum of 5 s per rating scale. Participants were asked to rate the effort required to bring the event to mind (1 = very easy, 6 = very effortful), feelings of re-experiencing the event (1 = not at all, 6 = completely), recall of setting (1 = not at all, 6 = distinctly), and perspective (1 = saw event through my own eyes, 6 = saw myself from an external perspective) (Scales adapted from Addis, Wong, & Schacter, 2007; Arnold, McDermott, & Szpunar, 2011). Participants were instructed to rate the experience of remembering and not the event itself. Response options on the screen appeared in square boxes representing the response box used in the MEG scanner, with each box representing one of the 4 buttons for each hand (excluding the thumb of each finger). Participants completed practice trials outside of the MEG to familiarize themselves with the task before moving on to the test trials. See Figure 1.

**Figure 1.**
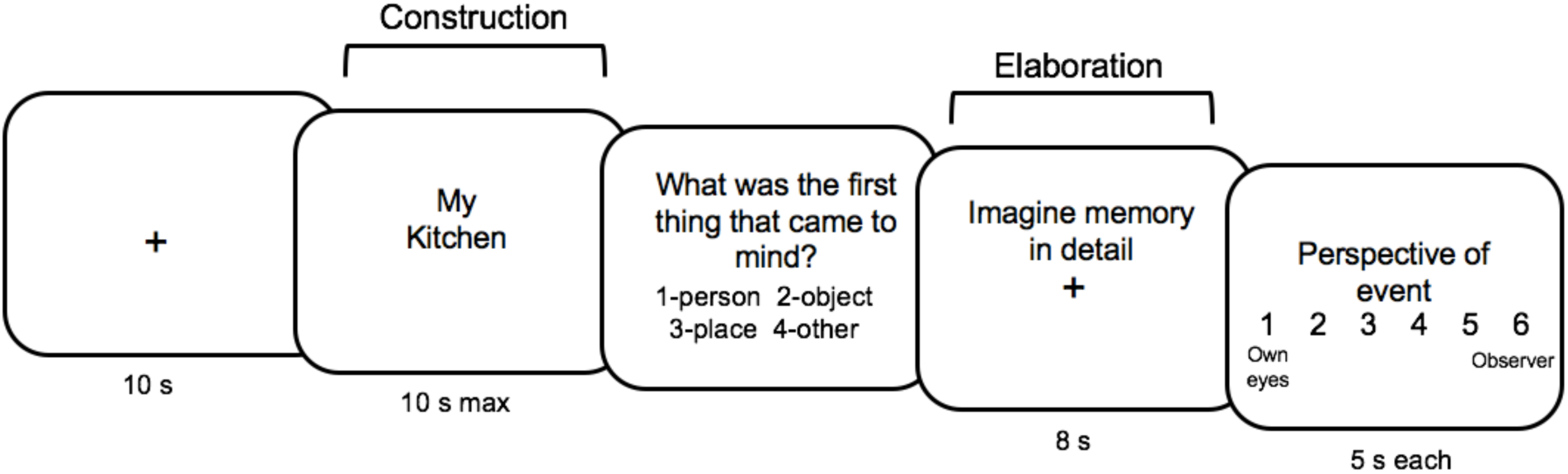
Autobiographical memory paradigm. Participants were cued with familiar locations and non-locations (people, objects), and told to recall a past event in relation to this word. They pressed a button when they had this event in mind.

#### 2.2.2. TMS procedure

Participants received cTBS to their left precuneus (MNI -14, -66, 56) and to a control region (vertex; MNI 0, -15, 74) on separate days. The order of these sessions was counterbalanced. Participants were blind to the type of stimulation they received (precuneus or vertex) and later interviews indicated they could not distinguish between the two. The precuneus was chosen as a target region based on our interest in this regions role in egocentric perspective during AM (Freton et al., 2014; Hebscher et al., 2018). We chose the left precuneus based on evidence showing that episodic/autobiographical memory is predominately associated with left-lateralized parietal activity (Kim, 2011; Rugg & Vilberg, 2013; Shimamura, 2011). The left precuneus target region was selected based on a custom meta-analysis of 13 studies with the keyword ‘egocentric’ using NeuroSynth (neurosynth.org). Within this map, the target region was selected by choosing the coordinates with the highest z-score that would be the most accessible with TMS (the most superficial area). Prior to stimulation, both stimulation site coordinates were warped from MNI to individual space and the stimulation site was chosen based on individual anatomy from whole-brain anatomical MRIs as the most superficial region that was closest to these coordinates.

At the beginning of the first stimulation session, resting motor threshold (RMT) was measured for each participant as the lowest intensity that produced motor evoked potentials (MEPs) above 50 µV in 5 out of 10 trials, recorded from the right first dorsal interosseous muscle. The Brainsight frameless stereotaxic neuronavigation system (Rogue Research, Montreal, Quebec, Canada) was used to target the selected stimulation sites. Four anatomical landmarks located on the face were used to co-register the anatomical MRI to the participant’s head. An infrared camera (Polaris Vicra, Northern Digital) recorded sensors attached to the participant and the TMS coil, allowing for real-time tracking of the TMS coil over the participant’s MRI. A biphasic Super-Rapid Stimulator 70-mm air-cooled Figure-8 coil (Magstim Co., Whitland, Dyfed, UK) was used to deliver a modified continuous theta burst stimulation (cTBS) at 80% RMT, lasting for approximately 40 s. The cTBS protocol consisted of 600 pulses arranged into bursts delivered every 5 Hz (200 ms), with each burst containing three pulses delivered at 30 Hz. The coil was positioned perpendicular to the stimulation site. Although standard cTBS protocols are delivered at 50 Hz, we decided to lower the burst frequency to 30 Hz due to limitations of the coil circuitry, leading to overheating at high intensities. Reducing the frequency of stimulation allowed us to stimulate at a higher intensity than would be possible at 50 Hz. Similar protocols have previously been shown to induce stronger MEP suppression compared to 50 Hz cTBS at a reduced intensity (Wu, Shahana, Huddleston, & Gilbert, 2012).

#### 2.2.3. MEG acquisition

MEG was recorded in a magnetically shielded room at the Rotman Research Institute with a 151-channel whole-head system with first order axial gradiometers (CTF MEG, Coquitlam, BC, Canada) (VSM MedTech Inc.), at a sampling rate of 625 Hz. Participants sat in an upright position and the behavioural task was projected onto a screen in front of them. For the first 4 participants MEG data was recorded continuously during the 30 minute behavioural task. We subsequently divided the behavioural task into three equal blocks approximately 10 minutes in length in order to reduce overall measures of head movement. To further minimize head movement, a towel was used to provide a tighter fit within the helmet. Head position was tracked at the beginning and end of each recording block by coils placed at three fiducial points on the head. Average head position across runs was used for source localization and was co-registered with fiducial points marked on the anatomical MRI. After acquisition, continuous signals were divided into epochs corresponding to each trial.

### 2.3. Behavioural analyses

To determine whether precuneus stimulation affects the behavioural dyanmics of AM retrieval, paired-samples t-tests were used to compare behavioural measures for precuneus and vertex stimulation sessions. Dependent variables were ease of recall ratings, reaction times, and place-other ratios (described below).

#### 2.3.1. Place-other ratio calculation

To measure the tendency to recall a location before other information, we calculated a ‘place-other’ ratio. A similar measure of early spatial recall is reported in Hebscher et al. (2018). We first excluded responses that matched the cue (e.g. location-cued memories where participants selected place as the first thing that came to mind). Each response type was then tallied across all other cue-types, allowing for calculation of likelihood ratios for each participant. To form a ratio of place responses versus other responses, we computed the proportion of place responses for person and object cues, the proportion of person responses for object and place cues, and the proportion of object responses for place and person cues. Because several participants had proportions of zero for either person or object responses (i.e. they never reported a person came to mind first when not cued with a person), we decided to sum these proportions to form an ‘other’, or non-location, responses condition. We then divided the proportion of place responses by the proportion of other responses to form a ‘place-other’ ratio. This ratio represented the likelihood of spontaneously reporting that a location first came to mind when not cued with one, relative to the likelihood of reporting that something else came to mind first. We were not able to compute place-other ratios for 1 participant for their precuneus stimulation session, and 1 participant for vertex stimulation session because these participants had proportions of zero for ‘other’ responses. Additionally, we were not able to record responses for the first thing that came to mind for one participant (vertex session) due to technical difficulties with the MEG response box. Thus, we had data for both sessions for a total of 20 participants.

### 2.4. MEG analyses

MEG data was preprocessed using Brain Electrical Source Analysis (BESA) Research 6.1. Noisy or bad channels were interpolated if they were central or defined as bad if in the periphery. We marked the same one channel as bad for 11 participants. One participant was found to have nosier data compared to others during their vertex stimulation session, and so 10 channels were defined as bad for this session. For each participant, we first performed a manual check of the data to remove large artifacts such as muscle tension. For periodic artifacts like eye-blinks, vertical and lateral eye movements and cardiac activity, we performed ICA removal, such that 1-3 components were defined for each participant based on their topography. After ICA correction, an automated artifact scan was run on all participant files to remove signal containing excessively high amplitudes that was not detected manually or through ICA correction. The data files were then averaged, yielding an average of 28.3 (SD = 3.0) (94.3%) accepted trials for construction and 27.6 (SD = 3.1) (92%) accepted trials for elaboration across participants. Next, the data was filtered using a low cut-off filter at 0.53 Hz (forward, 6dB/oct) and a high cut-off filter at 30 Hz (zero phase, 24dB/oct).

#### 2.4.1. ERF analyses: Effects of precuneus stimulation on retrieval dynamics

We performed paired-samples t-tests between the two stimulation sessions (precuneus vs vertex) to determine whether the precuneus plays a causal role in the temporal dynamics of retrieval. All ERF analyses were performed for 0-1000 ms of elaboration and construction stages using BESA Statistics 2.0, which includes a spatio-temporal permutation-based correction for multiple comparisons. We used a cluster alpha of 0.05, 1000 permutations, with clusters defined using a channel distance of 4 cm resulting in an average distance of 7.45 neighbors per channel.

#### 2.4.2. MEG source localization

We applied Classical Low Resolution Electromagnetic Tomography Analysis Recursively Applied (CLARA) to source localize significant results at the sensor level (BESA Research 6.1). CLARA iteratively localizes activity by reducing the source space during each LORETA estimation. Two iterations were computed with a voxel size of 7 mm^3^, and data were regularized using the default singular value decomposition cutoff of 1%. The solution was computed using an adult realistic head model and registered against the standardized BESA finite element model created from the average of 24 anatomical MRIs in Talairach-Tournoux coordinate space. We ran source localization on significant clusters as previously determined using paired t-tests and repeated measures ANOVAs in BESA Statistics. For each significant cluster, we computed source localization for a 30 ms time window around the peak time point.

#### 2.4.3. Late positive component

In a final set of analyses we sought to identify the LPC, a component commonly associated with recollection. We extracted amplitudes for the peak of this component (600-800 ms) over all right parietal sensors for memory elaboration. To determine if the LPC is associated with memory performance, we performed a series of correlations between peak LPC amplitude and subjective memory measures. Subjective memory measures included perspective rating, effort rating, place-other ratio, and memory vividness. Memory vividness was captured by the average of re-experiencing and setting ratings, which were positively correlated for both sessions (r > .50).

## 3. Results

### 3.1. Behavioural results

To determine whether precuneus plays a causal role in the behavioural dynamics of AM retrieval, we compared place-other ratios, reaction times, and ease of recall ratings between stimulation sessions. Paired samples t-tests revealed a non-significant but marginal effect of stimulation on place-other ratios (t(20) = 1.81, p = .085, CI [−.09, 1.34]), such that precuneus stimulation led to higher ratios compared to vertex stimulation (Figure 2a). This non-significant effect suggests that precuneus stimulation leads to a marginal increase in the tendency to recall locations before other information. Note that this analysis may have been underpowered due to missing place-other ratios for some participants. Average place-other ratios were 1.61 (SD = 1.71) for vertex stimulation sessions, and 2.18 (SD = 2.68) for precuneus stimulation sessions (Figure 2d). These findings show that participants are on average more likely to report that a location came to mind first, but show high variability in this tendency, consistent with previously reported findings (Hebscher et al., 2018).

**Figure 2.**
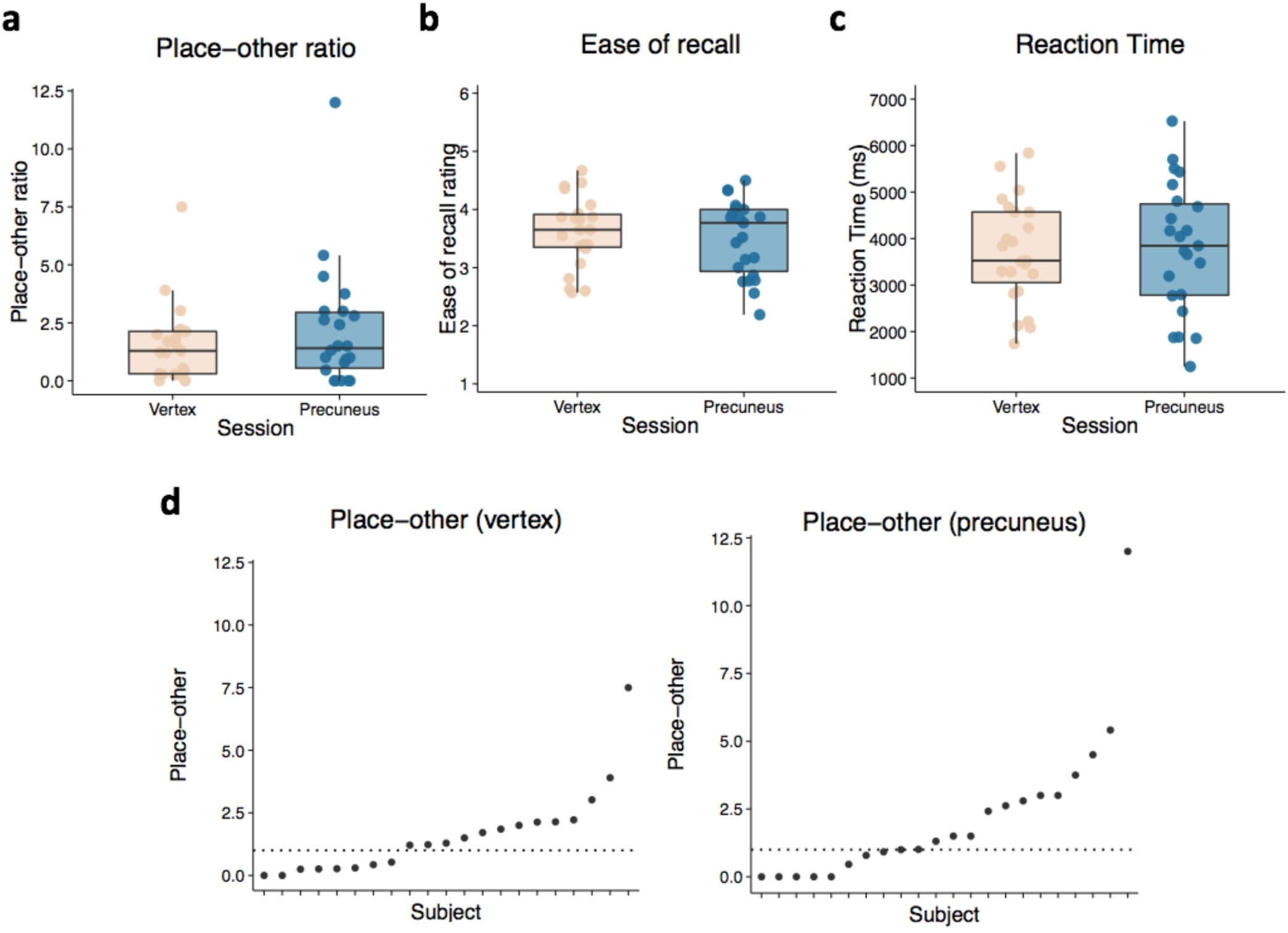
Behavioural results. (a) Paired-sample t-test showing non-significant comparisons between precuneus and vertex stimulation sessions on place-other ratio, (b) ease of recall rating, and (c) reaction time. (d) Distribution of place-other ratios for each participant. Dotted line reflects the point at which place and other responses were equal.

Precuneus stimulation did not significantly alter ease of recall ratings (t(22) = −0.76, p = .457, CI [−.41, .19]) or reaction times (t(22) = 0.37, p = .715, CI [−424, 609]). See Figure 2b-c. There were also no significant differences between perspective or vividness rating scales (all p’s > .46). An alternative analysis of these data is reported elsewhere, which shows an interaction between counterbalancing order and stimulation session on vividness ratings (Hebscher et al., 2019). This alternative analysis revealed that precuneus stimulation led to reduced vividness ratings when considering only the first session for each participant. Counterbalancing order did not interact with place-other ratios, ease of recall ratings, or reaction times, suggesting that there was no order effect on the behavioural dynamics of AM retrieval.

### 3.2. Evoked responses

Visual inspection of the evoked responses to cues during construction revealed typical, well-established components related to written word processing. Around 100 ms after cue onset there was a positive inflection in right occipital sensors resembling the P1 response, which reflects visual word processing (Kutas, Van Petten, & Besson, 1988). We also observed a well-formed M170 component over occipital and frontal sensors, peaking at around 100-200 ms post cue onset. The M170 corresponds with the N170 in EEG, which is a typical response to stimuli like faces and words, thought to reflect structural analysis of visual stimuli such as orthography for words (Bentin, Mouchetant-Rostaing, Giard, Echallier, & Pernier, 1999). We also observed a M400 (N400) over posterior sensors at around 350-500 ms, which typically reflects higher-level semantic processing of words (Bentin et al., 1999). Thus, cues elicited evoked responses consistent with well-established word processing components.

#### 3.2.1. Effect of precuneus stimulation on the neural dynamics of retrieval

We assessed the effect of precuneus stimulation on the dynamics of retrieval to determine the causal role of the precuneus in AM retrieval. Cluster-corrected paired-samples t-tests revealed that precuneus stimulation altered evoked activity at different stages of memory retrieval. During early construction, precuneus stimulation led to reduced negative amplitudes compared to vertex stimulation around midline frontal sensors, with precuneus stimulation appearing to delay the change in amplitude (cluster 1; 448-547 ms; p = .039; Figure 3a, left). Source localization for this effect (474-504 ms) failed to reach significance, but showed a trend towards reduced activity for precuneus stimulation in bilateral precuneus (p = .070). During later construction also around frontal sensors, we found that precuneus stimulation led to greater negative activity compared to vertex stimulation (cluster 2; 724-816 ms; p = .004; Figure 3a, right). Source localization for this effect (755-785 ms) identified a significant cluster in the right precuneus and premotor area with greater amplitudes for precuneus compared to vertex stimulation (p = 0.017) (Figure 3b). See Table 1a for cluster statistics. In both cases the more positive and negative deflections of the stimulation-related waveform appears to reflect delayed evoked responses, likely after the cue meaning had been processed and event construction had begun. These findings suggest that precuneus stimulation alters evoked responses during early AM construction such that an initial reduction or shift in activity is followed by an increase in activity, with both effects possibly arising from the precuneus.

**Table 1.**
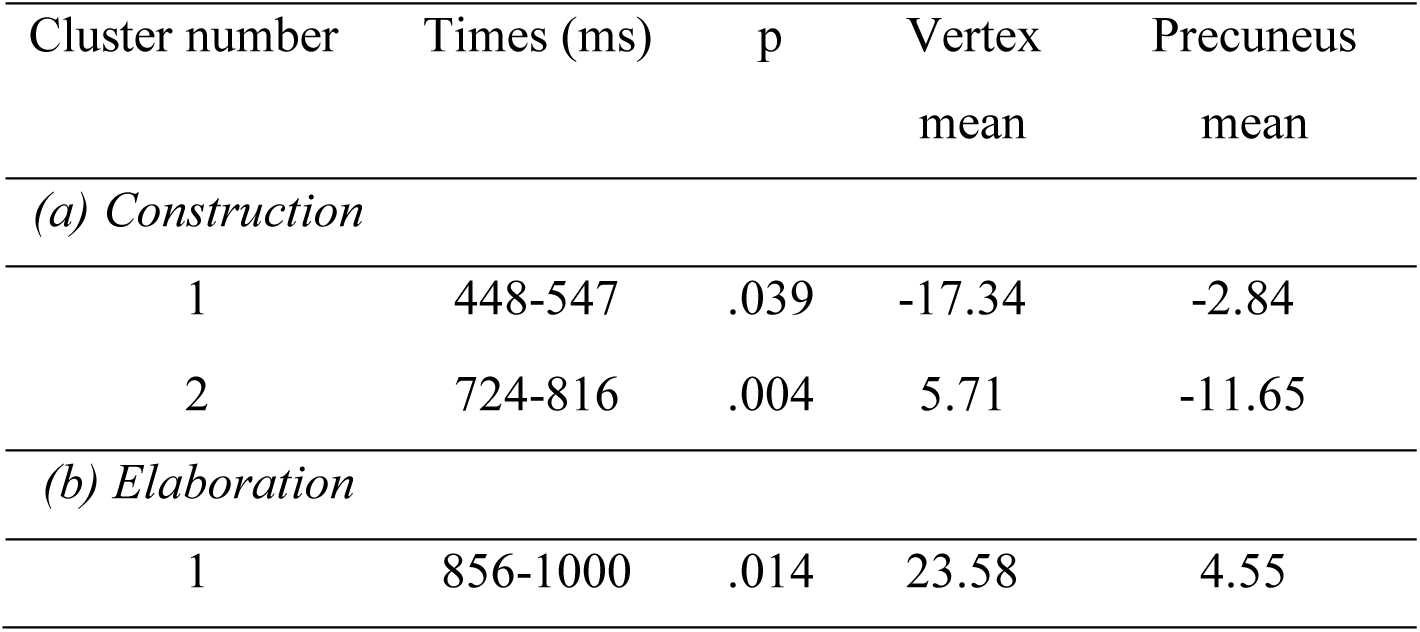
Cluster statistics for paired-samples t-tests

**Figure 3.**
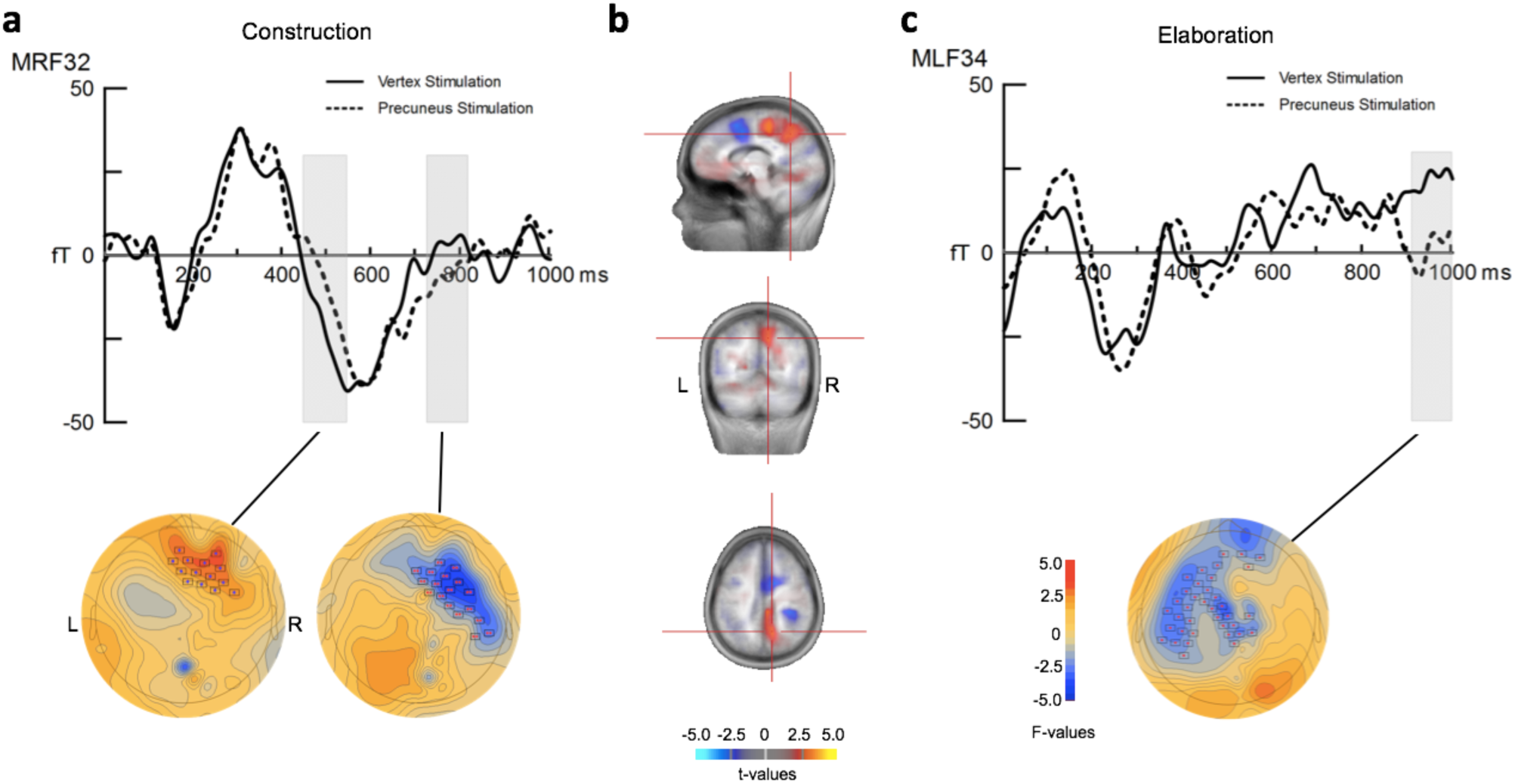
Differences between precuneus and vertex stimulation sessions during (a) construction and (c) elaboration. (a) ERFs for two significant clusters shown over a right frontal sensor with significant time points for that sensor highlighted in grey (top). Below, associated scalp maps show distribution of contributing sensors for both significant time intervals. (b) Source localization for the second significant cluster during construction (745-793). Warm colours show significantly greater right precuneus activation for precuneus compared to vertex stimulation, while cool colours are non-significant (p > .109). (c) ERFs over a left frontal sensor during elaboration with significant time points for that sensor highlighted in grey. Below, associated scalp map shows distribution of contributing sensors for significant time interval.

During later elaboration, amplitudes for memories following precuneus stimulation were again reduced compared to vertex stimulation over frontal-midline sensors (cluster 1; 856-1000 ms; p = .014; Figure 3c). Source localization for this effect was non-significant. See Table 1b for cluster statistics.

Together these results suggest that precuneus stimulation alters the temporal dynamics of retrieval at both construction and elaboration stages. During early memory construction, precuneus stimulation appears to slow down evoked responses, perhaps indicating a slower or delayed access to personal memories, while later in construction stimulation leads to more prolonged negative activity in the right precuneus. Precuneus stimulation also leads to a large reduction in amplitude during elaboration across a widespread cluster of frontal/midline sensors.

#### 3.2.2. Late positive component

In a final set of analyses, we sought to identify a late positive component commonly associated with recollection. The LPC is typically largest over posterior electrodes and is thought to emerge in parietal areas in EEG studies, with analogous positive modulations reported in MEG (Evans & Wilding, 2012; Tibon, Vakil, Levy, & Goldstein, 2014). Thus, we first visually examined posterior sensors for vertex stimulation sessions to determine whether AM retrieval similarly demonstrates the LPC. We identified a positive inflection resembling the LPC at approximately 600-800 ms over right parietal sensors during elaboration, the stage during which we expect recollection to occur. We did not observe this component during construction, likely because this stage is associated with search processes that typically differentiate AM from typical lab-based episodic memory tasks.

To determine if the LPC is associated with memory performance, we extracted amplitudes for the peak of this component (600-800 ms) for each participant for each right parietal sensor. Across the average of all right parietal sensors, there was a significant negative correlation between peak amplitude and perspective ratings such that greater amplitudes were associated with lower egocentric perspective ratings (r = −.54, CI [−.78, −.17], p = .008). See Figure 4a. Peak amplitude was not significantly correlated with memory vividness, effort, or place-other ratio (all p’s > .25). The same analysis for precuneus stimulation sessions revealed no significant correlation between LPC peak amplitude and perspective rating across all right parietal sensors (r = −.12, CI [−.51, .31], p = .583) (Figure 4b). Notably, the correlation between perspective rating and LPC peak amplitude was significantly greater for vertex stimulation than precuneus stimulation (z = −1.97, p = .025), indicating that this association is reduced or eliminated by precuneus stimulation. Paired-samples t-tests comparing peak amplitudes between sessions revealed a non-significant trend towards reduced LPC amplitudes for precuneus compared to vertex stimulation (t(22) = 1.96, CI [−.97, 24.7], p = .068).

**Figure 4.**
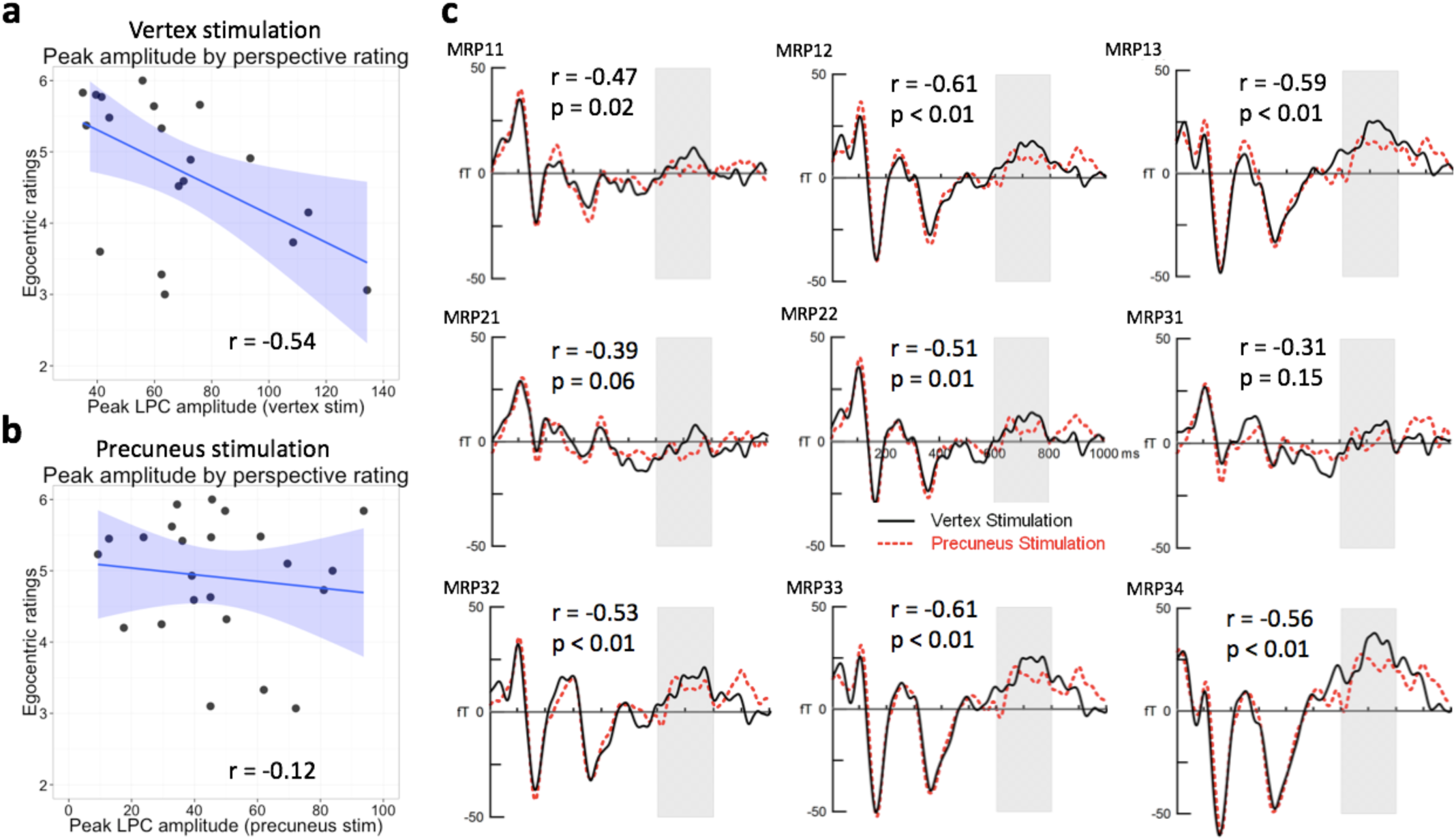
Late positive component. (a) Significant correlation between perspective rating and peak amplitude of the LPC across all right parietal sensors for vertex stimulation session, and (b) non-significant correlation for precuneus stimulation. Greater egocentric recall is reflected by higher values on the rating scale. (c) Plots for individual right parietal sensors for vertex (black) and precuneus (red) stimulation sessions. Corresponding *r* and *p* values for correlation with perspective ratings for vertex stimulation sessions are listed for each plot.

Next, we examined the correlation between perspective ratings and peak LPC amplitude for each right parietal sensor separately. For vertex stimulation sessions, the majority of sensors (7/9 sensors) were significantly correlated with perspective rating, while no sensors for precuneus stimulation sessions were significantly correlated with behaviour (all p’s > .256). As seen in Figure 4c, two sensors for vertex stimulation sessions were not significantly correlated with behaviour, and these sensors also demonstrated relatively small LPCs (MRP21, MRP31), while sensors with larger LPCs were more likely to be significantly and strongly correlated with behaviour. These findings show that greater LPC over parietal sensors is associated with the tendency to recall events from less of a purely egocentric perspective, likely reflecting a more flexible and variable experiential perspective. Furthermore, precuneus stimulation disrupts the relationship between LPC and perspective rating.

## 4. Discussion

The primary aim of this study was to elucidate the temporal dynamics of AM at the neural level. Based on the proposed role of the precuneus in the early dynamics of memory retrieval (Byrne et al., 2007), we set out to test the causal role of this region. Precuneus stimulation altered activity during early memory retrieval, delaying the evoked response in time, suggesting that this region plays a causal role in the neural temporal dynamics that support autobiographical memory retrieval. We also identified a positive modulation over posterior parietal sensors between 600-800 ms post-elaboration onset, analogous to the LPC, a well-established neural correlate of lab-based episodic recollection. Finally, we demonstrated that the LPC was associated with spatial perspective during AM recollection. Precuneus stimulation disrupted this association, demonstrating that perturbing the precuneus can break down the relationship between early neural modulations and spatial aspects of AM re-experiencing. These findings further suggest an association between spatial perspective and episodic recollection, in line with previous studies arguing that spatial context and episodic memory are importantly linked (Robin, 2018 for review).

### 4.1. Causal role of the precuneus in the dynamics of AM

We found that construction and elaboration stages of memory retrieval were differentially affected by precuneus stimulation. During early construction, precuneus stimulation led to reduced activity compared to vertex stimulation, reflected by a delayed rise of waveform amplitudes. This suggests that precuneus stimulation caused a delay or slowing of the evoked response during early construction, a stage during which participants are searching for and reconstructing a memory. Source localization for this effect revealed a non-significant trend towards reduced amplitudes in bilateral precuneus. Slightly later during construction, precuneus stimulation led to more negative amplitudes, which was source localized to the right precuneus. It is interesting that we stimulated the left precuneus and this later finding was localized to the right precuneus. One interpretation of these results is that precuneus stimulation initially delayed the evoked response to a cue by suppressing precuneus activity, but following this inhibition or delay the right precuneus compensated by increasing its activation, leading to a successfully retrieved memory. We also found that precuneus stimulation led to a large reduction in activity during later elaboration. Rather than reflecting a shift in activity, visual inspection of this finding suggested an overall inhibition of activity at this time point, which fits more with the inhibitory role of cTBS.

Space-based models of memory implicate a network of regions including the hippocampus, retrosplenial cortex, and precuneus in early space-based functions believed to be central to memory retrieval (Hassabis, Kumaran, & Maguire, 2007; Irish et al., 2015; Mullally, Vargha-Khadem, & Maguire, 2014; Becker & Burgess, 2001; Byrne, Becker, & Burgess, 2007). Our findings support these models, showing that the precuneus is causally involved in early memory processes, as early as 450 ms after cue presentation. It will be important for future studies to clarify the roles of the hippocampus and retrosplenial cortex in early memory processes to properly determine the validity of these space-based models of memory.

### 4.2. Late positive component

We aimed to identify the late positive component in our data and determine its relation to subjective aspects of AM. We identified a positive component resembling the LPC at approximately 600-800 ms during elaboration over right parietal sensors which was correlated with spatial perspective ratings. Precuneus stimulation eliminated this relationship, demonstrating a causal role for this region in the association between LPC peak and perspective rating. Interestingly, the significant relationship between LPC and perspective was such that participants with greater LPC amplitudes tended to recall events from less of a predominately first-person perspective. Perspective rating was a continuous scale which allowed participants to select the degree of first-person or third-person remembering, with mid-range ratings reflecting a combination of both perspectives. Notably, participants who scored the lowest on the perspective rating scale fell somewhere around the middle of the scale, suggesting that they were more likely to remember events from a combination of perspectives rather than from predominately third-person or first-person perspectives. Thus, mid-range ratings (associated with higher LPC amplitudes) may reflect more flexibility in the ability to translate perspectives, while higher ratings (associated with lower LPC amplitudes) may reflect less flexibility. Our results therefore suggest that participants more likely to flexibly shift perspectives during retrieval have greater peak LPC amplitudes. This finding fits with one previous study showing that parietal activity, in particular the central precuneus and right angular gyrus, is related to shifts in perspective during autobiographical memory retrieval (St. Jacques, Szpunar, & Schacter, 2016). It is also consistent with computational model predictions that translation between perspectives, or spatial updating, occurs repeatedly throughout memory retrieval (Bicanski & Burgess, 2018; Byrne et al., 2007). Thus, these data suggest that the LPC is important for the ability to translate spatial perspectives during memory recollection. Following precuneus stimulation, the LPC may no longer be important for or sensitive to perspective, perhaps because an important node in this translation circuit has been altered.

The LPC is typically characterized by a positive modulation occurring around 500-800 ms predominately over posterior sites, often but not always exhibiting a left-sided maximum (Rugg & Curran, 2007). This component has mainly been studied in the context of episodic memory where it has been consistently linked to recollection (Rugg & Curran, 2007). While the majority of these studies used EEG, analogous positive modulations have been reported in MEG (Evans & Wilding, 2012; Tibon et al., 2014). The LPC is sensitive to the amount of information recollected, suggesting that it reflects the representation or maintenance of recollected information (Vilberg, Moosavi, & Rugg, 2006; Wilding, 2000). Others have argued that this effect indexes attentional orienting to recollected information (Wagner, Shannon, Kahn, & Buckner, 2005). Our findings demonstrate that the LPC is also sensitive to perspective ratings, suggesting that spatial perspective affects some aspect of AM recollection. A previous study demonstrated the absence of LPC in a lab-based memory recognition paradigm in individuals with deficient AM (Palombo, Alain, Söderlund, Khuu, & Levine, 2015). Our findings demonstrate the direct presence of LPC during AM retrieval and importantly link this component to subjective aspects of recollection. Future studies are needed to further clarify the role of the LPC in autobiographical memory and its relationship to spatial perspective.

### 4.3. Conclusions

This study aimed to elucidate the temporal dynamics of AM retrieval, which few studies to date have done. We show that the precuneus plays a causal early role in the neural dynamics of retrieval, in line with the proposed role of this region (Byrne et al., 2007). We show also that an established neural component of recollection is related to the tendency to recollect AMs from mixed spatial perspectives and that the precuneus is casually involved in this relationship. These findings provide novel insights into the role that spatial information plays in the temporal dynamics of retrieval and help clarify the neural correlates of early memory retrieval.

## Acknowledgements

We would like to thank Brahm Sanger and Kyle Nealy for their assistance in testing participants, and Jed Meltzer for help with experimental design.

## Funding

This work was supported by the Natural Sciences and Engineering Research Council of Canada (grant 378291). Additional support was provided by the Natural Sciences and Engineering Research Council of Canada Postgraduate Scholarship

